# Profiling neurons surrounding subcellular-scale carbon fiber electrode tracts enables modeling recorded signals

**DOI:** 10.1101/2022.10.07.511373

**Authors:** Joseph G. Letner, Paras R. Patel, Jung-Chien Hsieh, Israel M. Smith Flores, Elena della Valle, Logan A. Walker, James D. Weiland, Cynthia A. Chestek, Dawen Cai

## Abstract

Characterizing the relationship between neuron spiking and the signals electrodes record is vital to defining the neural circuits driving brain function and informing computational modeling. However, electrode biocompatibility and precisely localizing neurons around the electrodes are critical to defining this relationship. Here, we show the ability to localize post-explant recording tips of subcellular-scale carbon fiber electrodes and surrounding neurons. Immunostaining of astrocyte, microglia, and neuron markers confirmed improved tissue health. While neurons near implants were stretched, their number and distribution were similar to control, suggesting that these minimally invasive electrodes demonstrate the potential to sample naturalistic neural populations. This motivated prediction of the spikes produced by neurons nearest to the electrodes using a model fit with recorded electrophysiology. These simulations show the first direct evidence that neuron placement in the immediate vicinity of the recording site influences how many spike clusters can be reliably identified by spike sorting.

## Introduction

Recording and interpreting neural activity in the mammalian cortex is paramount to understanding brain function. Capturing the signals of individual neurons yields the most fundamental dynamics of neural activity^1,2^, thereby providing the highest precision when decoding neural circuits^3^. As intracortical electrode recordings have a high spatiotemporal resolution to capture individual neurons’ signaling associated with fast-paced behaviors^4^, electrode architectures with many dense recording sites are desired for sampling large populations of neurons^1,5^.

Currently, the most widely used intracortical electrodes are composed of silicon shanks fabricated using standard cleanroom techniques^6^. These electrodes are becoming increasingly sophisticated, with newer designs approaching or exceeding a thousand densely-packed recording sites^7–11^. For example, the Neuropixels probe has demonstrated simultaneous recording from hundreds of individual neurons along its length^8,9,12^. Planar silicon electrode arrays, e.g., the Utah Electrode Array (UEA), can sample wide areas of cortex^13^ for use in BMIs^14^, enabling the restoration of function lost to neurological disease^15–17^. Moreover, the recent advancements of multi-shank Michigan-style electrodes^10^, such as the Neuropixels 2.0^9^, and variable-length UEA-style electrodes, such as the Sea of Electrodes Array^7^, signify that recording neuronal populations in 3D is possible.

Extensive evidence, however, indicates that chronic implantation of silicon electrodes can elicit a multi-faceted foreign body response (FBR) at the implant site, which includes significant glial scarring^18^, high microglia presence^19^, large voids of tissue^20,21^, blood-brain-barrier disruption^22^, and neurodegeneration^23,24^. Recent studies have uncovered more effects, including hypoxia and progressive neurite degeneration^25^, a shift toward inhibitory activity^26^, myelin injury and oligodendrocytes loss^27^, and mechanical distortion of neurons^28,29^. RNA sequencing of implant site tissue also yielded differential expression of more than 100 genes, signifying the FBR results in complex biological changes^30^. Moreover, recorded signal amplitudes degrade over chronic time periods^31,32^ and can both increase and decrease during experimental sessions^31,33^, which have been attributed to the FBR^21^. The chronic neuron loss, particularly within the single unit recording range^34,35^, brings to question silicon electrodes’ ability to reliably record activity attributed to individual neurons chronically. Silicon probes’ large size has been implicated as a primary reason for the FBR^19,36,37^. Many newer electrode technologies have been designed to overcome the FBR^1,36,38^. In particular, smaller electrodes that have cellular to subcellular dimensions^39–43^ generate smaller tissue displacement^44^ and reduced FBR and neuron loss^36,37^, suggesting an ability to record more naturalistic neural populations.

While computational models have been proposed to decode the relationship between recorded spikes and contributing neurons^45–49^, their validity remains unresolved without empirical measurements of neuron locations and their spike timing^50^. Attempts at acquiring these “ground truth” measurements have been performed using tetrodes or electrodes with similar geometry^34,47,51^, which Marques-Smith *et al*.^50^ assert as differing considerably from state-of-the-art electrodes such as Neuropixels^8,50^, under *ex vivo* conditions^52,53^, simulated data^54–56^, or acutely *in vivo*^57,58^ and with low resolution in cell localization^50^. As modeling suggests that biofouling in the electrode-tissue interface influences neural recording quality^49^, “ground-truth” must be measured in cases where electrodes are implanted *in vivo* over long time periods. Precisely localizing chronically implanted electrodes *in situ* and surrounding neurons has begun to bridge this gap^39,40,59,60^.

Subcellular-scale (6.8 μm diameter) carbon fiber electrodes that elicit a minimal FBR and maintain high neuron densities after chronic implantation^43,59,61,62^ are ideal candidates for acquiring these “ground truth” recordings. Previously, we developed a “slice-in-place” technique to retain the electrodes in brain slices for localizing the recording tips in deep brain structures^59^. However, brain curvature and the bone screws required for skull-mounted headcaps render this method incompatible with fibers implanted in shallower cortical regions. In this report, we demonstrate that explanting carbon fiber electrodes from cortical recording sites followed by slicing thick horizontal brain sections with headcaps removed retains the ability to localize the tips with high-resolution 3D images. We used rats chronically implanted in layer V motor cortex to assess neuronal health and glial responses and for modeling the relationship between recorded spikes and surrounding neurons. From 3D reconstructions of neuron somas, we found almost no loss in neuron count per volume, although neurons were stretched compared to neurons in the contralateral hemisphere. The distance of the nearest neuron to implanted fibers was similar to that of simulated electrodes positioned in the contralateral hemisphere. Given the minimal disruption in surrounding neurons, we modeled the extracellular spikes that could be recorded from the neuron population at the implant site, which suggested that their natural distribution is a fundamental limiting factor in the number of spike clusters that can be sorted.

## Results

### Assessment of foreign body responses induced by whole carbon fiber electrode arrays implanted chronically

We first verified that subcellular-scale (6.8 μm diameter) carbon fiber electrode arrays implanted for this study yielded minimal FBRs similar to those observed with our other carbon fiber electrode designs^59,61,62^. To assess the FBR along the arrays, we implanted one High Density Carbon Fiber (HDCF)^63^ electrode array targeting layer V motor cortex in each of three rats for 12+ weeks. This design included tapering silicon support tines that extended up to the last 300 μm of the fibers’ length to enable direct insertion (Figure 1A)^63^. Similar to other implanted silicon electrodes^18,38^, we found elevated GFAP staining around the silicon supports, indicating astrocyte enrichment as part of the FBR and signifying the implant site (Figure 1B). The explanted electrodes left an approximately straight row of black holes in each image plane, illustrating the electrode tracts in 3D in brain slices containing the tips. At depths near putative electrode tips and far from the silicon supports (~300 μm), while enriched astrocytes persisted, minimal microglia responses and high neuron densities were observed (Figure 1C and Video 1). These microglial and neuronal responses appeared similar to those observed in our previous reports^59,61,62^ and those located in symmetrical regions in the contralateral hemisphere (Figure 1D). It is worth noting that we observed varying degrees of FBRs in histology showing the implant sites for all three rats (Figure 1 and Figure S1). Such variation may reflect differing structural damage during implantation^64^. Furthermore, we quantified glial fluorescent intensities with known carbon fiber locations for the first time. GFAP and IBA1 intensities were elevated up to 200 μm and 60 μm from putative electrode tips (Figure S2), respectively, which is considerably closer than our previous measurements with silicon shanks^61^.

**Figure 1.**
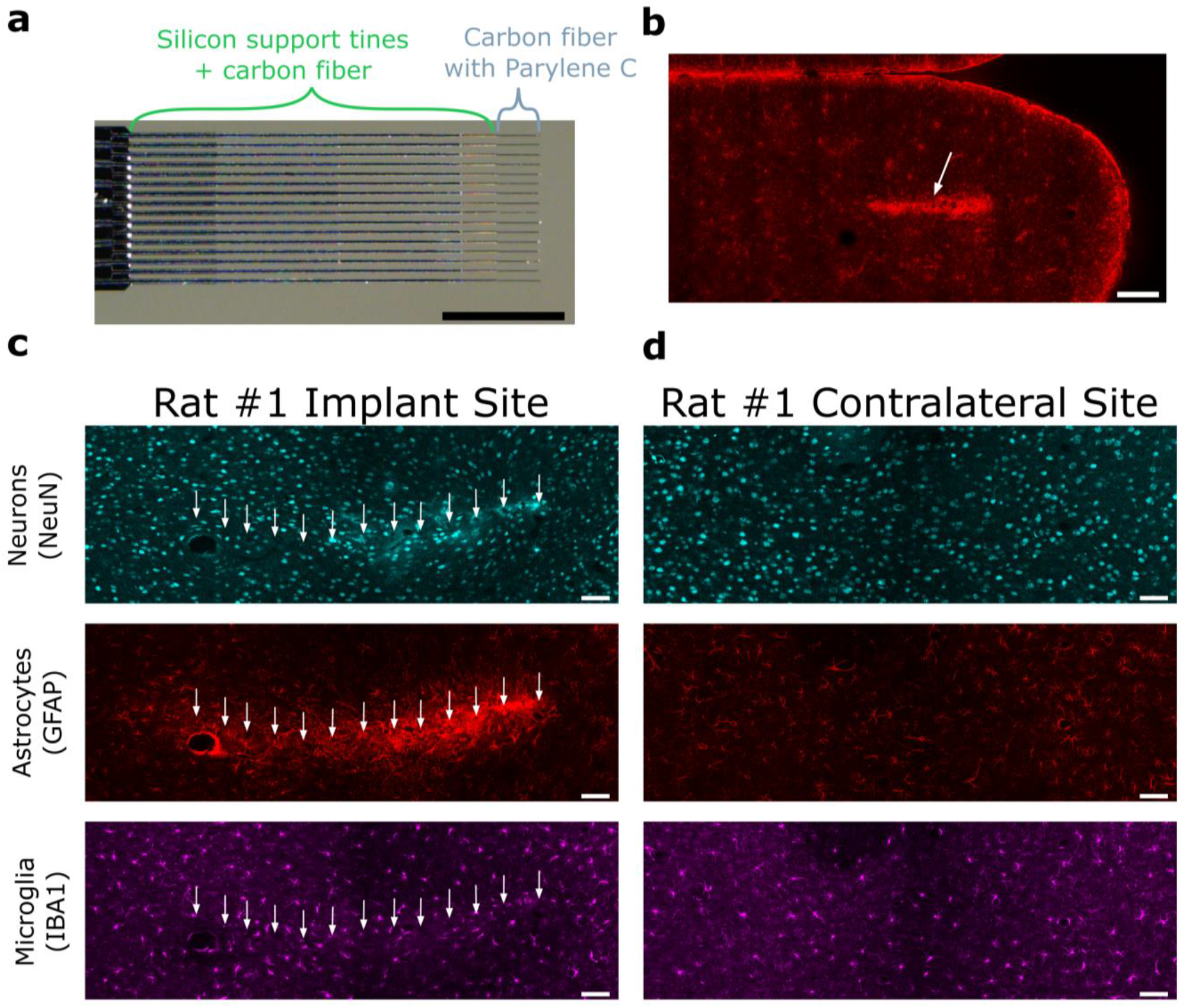
Electrode array identification and assessment of foreign body responses induced by whole carbon fiber electrode arrays implanted chronically. (A) Example of a HDCF electrode array implanted for this study. Tapering silicon support tines supported the fiber until the last 300 μm of the fiber length, where the tip was the recording site. Scale bar, 1 mm. Image was slightly rotated in ImageJ. (B) Confocal image of astrocytes (GFAP) in the brain slice immediately dorsal to the brain slice with tips for rat #2. Glial responses dorsal to the tips from the support tines helped localize electrode tracts in ventral slices with tips. The arrow points to the implant site. Scale bar, 500 μm. (C) Representative histology of the implant site in rat #1 (layer V motor cortex). Imaging plane was estimated to be up to 42 μm ventral of some tips, and up to 12 μm dorsal to others.(D) Representative histology of a site contralateral to the implant site for rat #1 with similar stereotaxic coordinates. Imaging settings were the same. (C and D) Scale bar, 80 μm. (B-D),Cyan: NeuN (neurons); red: GFAP (astrocytes); magenta: IBA1 (microglia). White arrows point to fiber tracts identified in the tissue. Images were stitched^65^ and contrast adjusted in ImageJ. See also Figure S1 for whole-array histology of rats #2 and #3.

### Carbon fiber recording sites can be localized in post-explant motor cortex with cellular confidence

Previously, we localized the recording site tips of similar carbon fiber electrodes in deep brain regions using our “slice-in-place” method^59^. However, using this method to cryosection implants at superficial brain regions, such as motor cortex, is difficult due to the intrusion of supporting skull screws and the curvature of the brain. Here, we explanted the electrodes and estimated tip locations using biomarkers and visual cues captured with submicron-resolution confocal imaging instead. Tips could be identified by a combination of factors, including the FBR itself. Elevated GFAP immunostaining typically delimited electrode tracts as astrocytes enriched around a series of dark holes, particularly at shallower depths (Figure 1B), but also at depths close to some tips. Simultaneous transmitted-light imaging demonstrated that these holes were not immunostaining artifacts (Figure 2A). Such dark holes also appeared in other fluorescent channels contrasting with background staining. Successive confocal imaging produced high-resolution and high-contrast renditions in 3D. Collectively, we utilized the disappearance of these holes in the dorsal-ventral direction to corroborate fiber tracts and localize putative tips. For instance, as shown in Figure 2B and Video 2, at the dorsal side of the brain slice, GFAP+ astrocytes wrapped around the dark hole of the electrode tract, signifying the fiber’s pre-explant location. This hole could then be followed in the ventral direction to a depth where it was rapidly filled in by surrounding parenchyma and background fluorescence, signifying the electrode tip.

**Figure 2.**
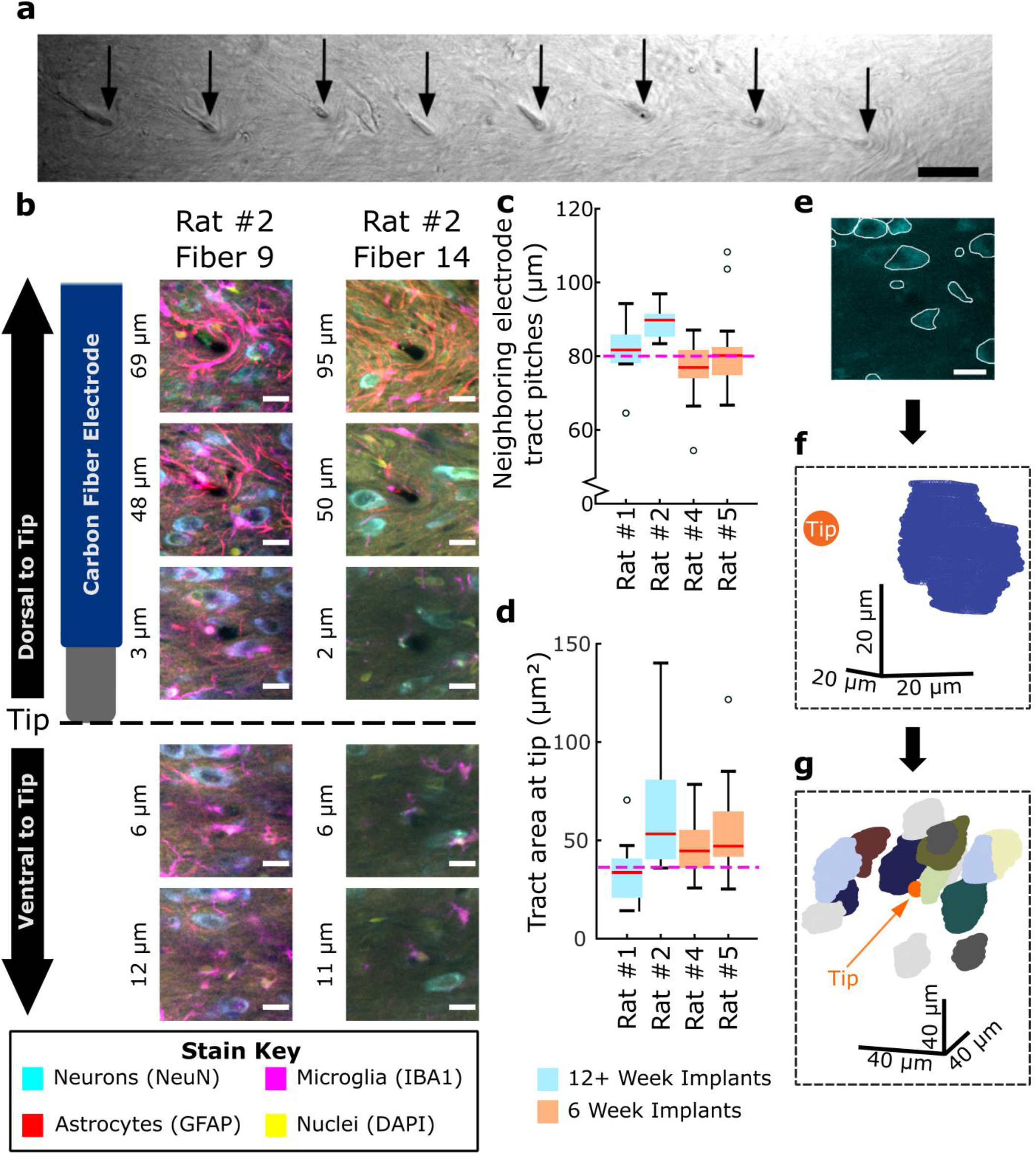
Carbon fiber recording sites can be localized in post-explant motor cortex with cellular confidence. (A) Transmitted light image of 8 fiber tracts identified in rat #2. Black arrows point to dark holes where fibers had been implanted. Scale bar, 50 μm. (B) Recording site tips of carbon fiber electrodes were identified by following the electrode tracts in the dorsal to ventral direction in 3D imaging volumes. Localization of two representative fibers is shown, with cartoon electrode accompanying (left). A black hole surrounded by glia or a disturbance in tissue, or both, is easily identified dorsal to the electrode tip (top). The tract was followed in the ventral direction until the hole was rapidly filled in by surrounding parenchyma and background fluorescence (images below dotted line). Scale bar, 20 μm. Color scheme is the same as in Figure 1, with yellow for cellular nuclei (DAPI) added. Images were contrast adjusted, histogram matched^66^ (twice for DAPI), and 3D Gaussian filtered^67^ in ImageJ for easier visualization. (C) Electrode pitches measured for implanted fibers (N=57 pitches). Dashed line is the expected value (80 μm) from the design. (D),Cross-sectional area of fiber tracts at the tips (N=61 fibers). Dashed line is the expected hole area for an uncoated fiber (36.3 μm^2^). (C and D) Rats implanted for 12+ weeks shown in blue (N=2), rats implanted for 6 weeks shown in orange (N=2). e-g, Quantification of neurons surrounding the recording site tips through neuron tracing. (E) Representative z-step (imaging plane) of neurons collected with confocal microscopy. The white outlines are traces that follow the edge of each neuron captured in the z-step imaging plane. Image is contrast adjusted, histogram matched, and median filtered. Scale bar, 20 μm. (F) Tracing the neuron in each z-step that it was captured yielded a 3D scaffold of points (blue points). (H) Tracing neurons with centroids within 50 μm from the recording site tip center yields an electrode-specific model of the surrounding neurons.

We localized 29 putative tips of 31 fibers that were implanted for 12+ weeks (N=2 rats), and 32 of 32 tips in an additional cohort implanted for 6 weeks (N=2 rats). Two tips in rat #1 were not found, as their locations were likely included in a slice where some electrode tracts merged (data not shown). The distance between adjacent putative electrode tracts was measured at 82.1±9.2 μm, (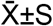, N=57 pitches), which was close to the array design pitch (80 μm) (Figure 2C). The cross-sectional area of the tips was 50.5±24.7 μm^2^ (N=61 fibers) and comparable to the expected 36.3 μm^2^ from a bare carbon fiber (Figure 2D). These measurements indicated that we correctly identified fiber locations. To estimate the precision in localizing tips, three observers independently corroborated tip locations (Table 1). The median absolute difference between estimates was 9.6 μm, suggesting subcellular precision. However, this difference was considerably lower in rats implanted for six weeks (5.2 μm) than 12 (14.7 μm), likely the result of better tip positioning and elevated background staining. Furthermore, that the median difference in the horizontal plane was 2.2 μm suggests that tip depth contributed more to tip localization error than tract position.

**Table 1.** Summary of comparisons of fiber tip localization by three reviewers. Each of three reviewers trained in histology localized putative carbon fiber tips and the absolute Euclidean distances between the localizations for each fiber were measured for comparison. Therefore, the number of comparisons was three multiplied by the number of fibers. Summary statistics were taken from the full samples of comparisons. The difference in the A/P and M/L plane was measured because images were collected in horizontally sliced brain sections, while the D/V axis was the axis of z-stack imaging. A/P: anterior/posterior. M/L: medial/lateral. D/V: dorsal/ventral.

| Rat(s) | Number of Fibers | Median Absolute Difference ($\mu\text{m}$) | Interquartile Range ($\mu\text{m}$) | Median Absolute Difference in A/P & M/L ($\mu\text{m}$) | Median Absolute Difference in D/V ($\mu\text{m}$) |
| --- | --- | --- | --- | --- | --- |
| Rat # 1 (12 Weeks) | 13 | 13.7 | 20.4 | 5.8 | 12.0 |
| Rat #2 (12 Weeks) | 16 | 15.4 | 16.6 | 1.8 | 15.0 |
| Rat # 4 (6 Weeks) | 16 | 6.9 | 9.2 | 1.4 | 6.6 |
| Rat #5 (6 Weeks) | 13 (3 excl.) | 4.3 | 6.7 | 2.7 | 2.4 |
| 12 Week Rats | 29 | 14.7 | 17.5 | 2.9 | 13.8 |
| 6 Week Rats | 29 | 5.2 | 9.0 | 1.8 | 4.8 |
| Pooled | 58 | 9.6 | 15 | 2.2 | 8.7 |

### Neuron soma morphology is geometrically altered surrounding carbon fiber tips

Having localized the recording site tips with high confidence and captured nearby neurons with submicron resolution imaging, we sought to assess changes in neuron soma morphology near the implants, as previous studies reported that neurons surrounding silicon electrode implants were mechanically stretched compared to neurons in non-implanted regions^28,29^. We traced the 3D outlines of nearby neuron somas (N=945 neurons, N=28 fibers) and somas in symmetric contralateral sites (N=1125 neurons) from NeuN staining images (Figures 2E-G) to measure soma shape and volume within a 50 μm radial sphere from implanted fiber tips and hypothetical tips positioned in the contralateral hemisphere. Here, we also found that neurons close to the fiber tips appeared to be stretched in one direction compared to neurons in contralateral tissue (Figures 3A and 3B). To quantify the degree of soma distortion, the cell shape strain index (CSSI) was calculated using Equation 1^28,29^. The CSSI was defined as the ratio of the difference to the sum of the longest axis and the shortest axis of the soma, where a high CSSI indicates the soma has a more asymmetric shape and a CSSI near zero indicates a nearly circular soma^28^. In Figure 3C, we plot the distributions of neuron soma CSSIs within a 50 μm radius from implanted fibers (0.53±0.23, 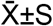, N=349 neurons), and found those were 112% larger than those at the contralateral site (0.25±0.12, 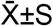, N=1125 neurons). Plotting the CSSI over the distance of each neuron relative to carbon fiber tips (red dots) allowed us to fit a linear regression line (red line) to extrapolate the distance at which neurons at the implant site would have CSSIs similar to the average CSSI (blue line) measured for neurons in the contralateral hemisphere (Figure 3D). The slope of the linear regression (R=0.15, p=5×10^−3^) predicted this distance is 147 μm. Since there was a morphological change for neurons in the close vicinity of the electrodes, we were also curious whether there was a concomitant change in neuron soma volume, which has not been investigated in previous FBR studies to our knowledge. We found that neuron somas within 50 μm from fibers were 23% smaller (2.3×10^3^±1.5×10^3^ μm^3^, N=349 neurons) than those in the contralateral hemisphere (2.9×10^3^±1.3×10^3^ μm^3^, N=1125 neurons) (p<0.001, Kolmogorov-Smirnov Test, Figure 3E). Fitting soma volumes (red points) over distance to fibers resulted in a regression (redline, R=0.13, p=0.012) that predicted the neuron volume would increase to that of the contralateral hemisphere (blue line) at 72 μm away from the fibers (Figure 3F).

**Figure 3.**
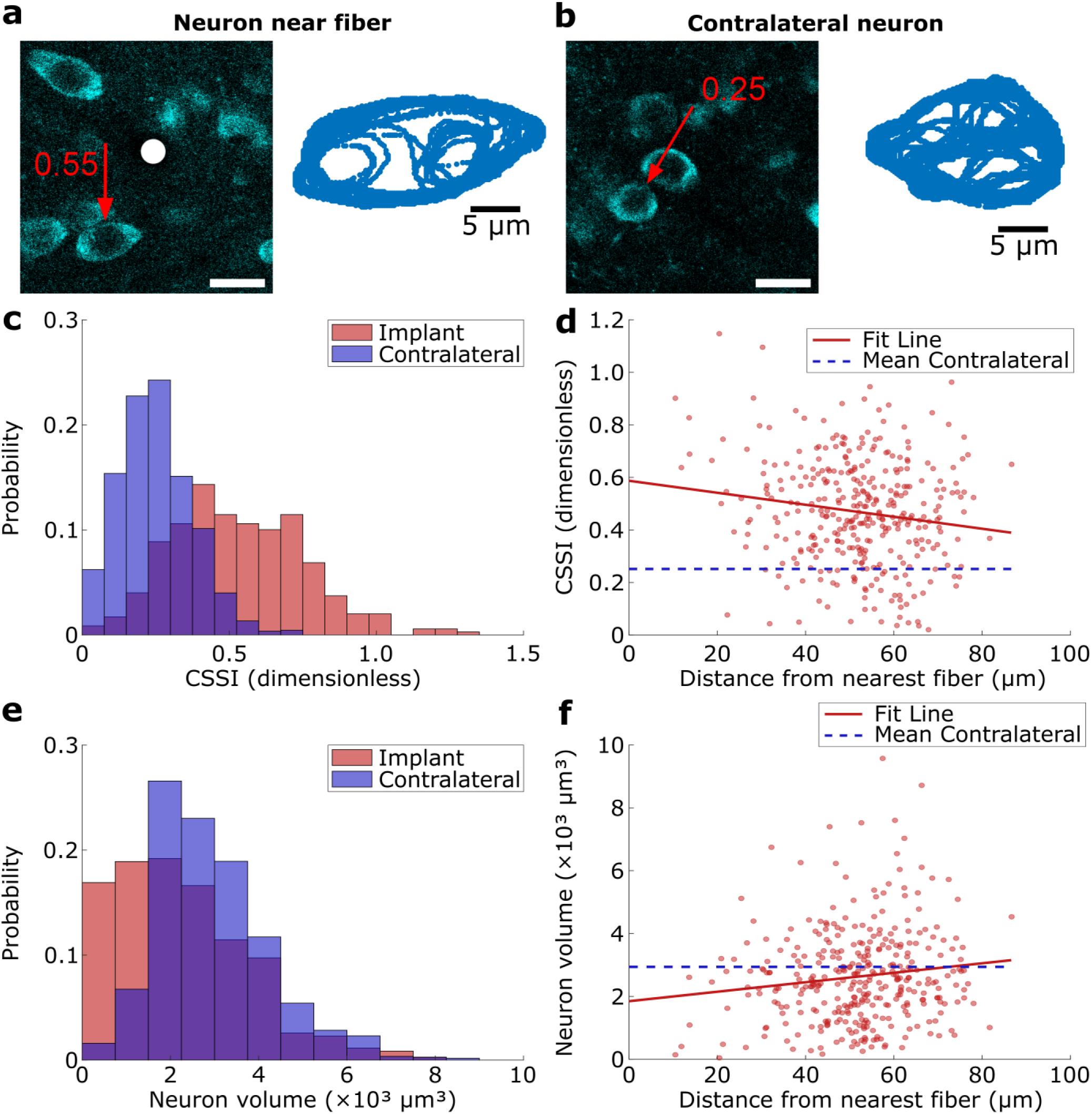
Neuron soma morphology is geometrically altered surrounding carbon fiber tips. (A) NeuN staining showing a representative neuron (CSSI^28^ = 0.55) stretched by the presence of an implanted fiber (white dot) (left) and corresponding neural scaffold (right). Scale bar (left), 20 μm. (B) Representative neuron in contralateral tissue (CSSI=0.25) (left) and corresponding neural scaffold (right). This neuron has a lower CSSI and is more circular. Scale bar (left), 20 μm. (C) Histograms showing the CSSIs calculated for neurons located within 50 μm from recording site tips (N=349 neurons) and neurons in contralateral tissue (N=1125 neurons) without implants (N = 2 rats, 28 fibers). CSSIs around fibers were significantly larger (Kolmogorov-Smirnov test, p<0.001). (D) CSSI vs. neuron distance (N=355 neurons) from tip. Linear regression (solid line) yielded fit line y=−2.3×10^−3^x + 0.59 (R=0.15, p < 0.01). Mean CSSI (0.25) for contralateral neurons shown for comparison (dashed line). (E) Same as (C), but with neuron somal volumes (N=349 neurons around fibers, N=1125 neurons in contralateral). Distributions are significantly different (Kolmogorov-Smirnov test p<0.001). f(F) Same as (D), but with neuron somal volumes (N=355 neurons). Linear regression (solid line) yielded fit line y=15x + 1848 (R=0.13, p<0.05). Mean volume (2.9×10^3^ μm^3^) for contralateral neurons is shown for comparison (dashed line).

### Neuron placement around implanted carbon fibers resembles naturalistic neuron distributions

As we observed many neurons surrounding the arrays, we desired to quantify how naturally these neurons were distributed in the fibers’ immediate vicinities. Using the centroids of the aforementioned 3D reconstructions of neuron somas (Figures 2E-G), we quantified neuron densities within a 50 μm radial sphere from implanted and hypothetical control fibers. The neuron density around implanted tips was 3.5×10^4^±0.9×10^4^ neurons/mm^3^ (N=16 fibers), which was 82±22% of that measured around hypothetical tips (N=2932 fibers). In contrast, conventional single-shank silicon electrodes retain 40% of a healthy neuron density within 50 μm^24^ and 60% within 100 μm^23^. As spikes from single neurons can putatively be separated into individual clusters within 50 μm from an electrode^34,35^, the higher density around carbon fibers within this range may explain their improved recorded unit yield over silicon probes^61^.

Having confirmed that carbon fibers preserve most neurons within 50 μm, we examined whether the nearest neuron positions relative to the tips were altered. This is important for modeling because the extracellular spike amplitudes recorded is inversely proportional to the distance between neurons and recording sites^5,68,69^. To do so, we plot histograms of the distances between the nearest six somas’ centroids to each electrode tip (Figure 4A) and compared them to the distances to each hypothetically positioned probe in the contralateral hemisphere (Figure 4B). Importantly, the neuron nearest to tips was measured at 16.8±4.5 μm (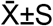, N=28 fibers) away, which was only 0.6 μm and not significantly farther away (p=0.55, two sample t-test) than the neuron nearest to hypothetical tips in the contralateral hemisphere (16.2±4.8 μm, N=2932 simulated fibers). We summarized the nearest six neuron positions in Table 2. Of those six positions, the difference between neurons of matching positions around implanted and hypothetical fibers was of subcellular scale at 2.6±1.1 μm 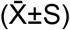. These results suggest that the neurons, which were mostly preserved with approximately proper positions, surrounding carbon fiber electrodes may have produced naturalistic physiological spiking activity.

**Table 2.** Nearest neuron positions surrounding carbon fibers. Closest six neuron positions relative to implanted carbon fiber electrodes (implant) and hypothetical fibers placed in contralateral tissue (contralateral). The number of fibers is differen t between positions because some fiber tips were too close to the top of the brain section to measure subsequent neuron positions (see Methods). The difference column shows difference in relative Euclidean distance between implanted fibers and hypothetical fibers in the contralateral hemisphere for each position.

| Neuron Position | Distance (Implant)<br>$\bar{X} \pm S \text{ } \mu\text{m}$ | Number of Fibers | Distance (Contralateral)<br>$\bar{X} \pm S \text{ } \mu\text{m}$ | Number of Simulated Fibers | Difference ($\mu\text{m}$) | Significance (p value) |
| --- | --- | --- | --- | --- | --- | --- |
| 1<sup>st</sup> | $16.8 \pm 4.5$ | 28 | $16.2 \pm 4.8$ | 2932 | 0.6 | 0.55 |
| 2<sup>nd</sup> | $23.2 \pm 6.1$ | 25 | $21.2 \pm 4.5$ | 2932 | 2.0 | $2.6 \times 10^{-2}$ |
| 3<sup>rd</sup> | $27.7 \pm 6.0$ | 23 | $24.7 \pm 4.2$ | 2932 | 3.1 | $4.0 \times 10^{-4}$ |
| 4<sup>th</sup> | $30.5 \pm 4.5$ | 22 | $27.4 \pm 3.9$ | 2932 | 3.2 | $1.8 \times 10^{-4}$ |
| 5<sup>th</sup> | $33.0 \pm 4.8$ | 22 | $29.6 \pm 3.7$ | 2932 | 3.4 | $2.8 \times 10^{-5}$ |
| 6<sup>th</sup> | $35.1 \pm 4.4$ | 22 | $31.7 \pm 3.6$ | 2932 | 3.4 | $8.5 \times 10^{-6}$ |

**Figure 4.**
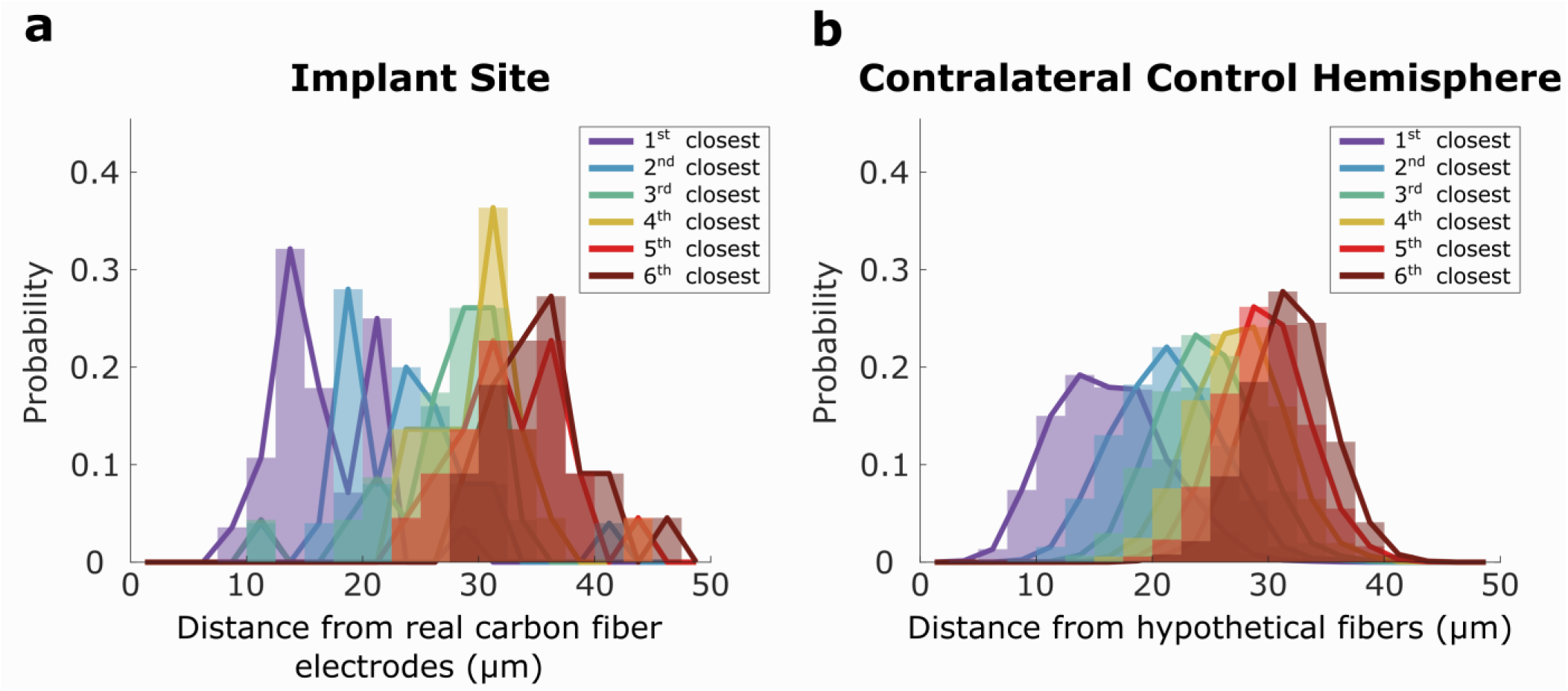
Neuron placement around implanted carbon fibers resembles naturalistic neuron distributions. (A) Histograms of the relative distances between the nearest six neurons and implanted carbon fiber electrodes in N=2 rats. (B) Histograms of the relative distances between the nearest six neurons and points chosen as hypothetical (faux fibers) fibers in the hemisphere contralateral to the implant sites in N=2 rats. The contralateral sites had similar stereotaxic coordinates to the implant sites and were within the same brain sections. (A and B) see Table 2 for a list of mean distances at each position.

### Modeling suggests neuron distribution contributes to the number of sorted spike clusters

Previous work suggests that spikes can be sorted into clusters from individual neurons that were recorded as far as 50 μm away from an electrode^34,35^. It has been assumed that each spike cluster includes action potentials from a single nearby neuron^35,70^, but it is possible that the electrode records similar spike waveforms generated from multiple neurons^71^. At the same time, it has been widely noted that the number of clusters determined by spike sorting, reported to be between one and four^72^ and between one and six clusters^35,54,73^, is at least an order of magnitude lower than the neuron count within a 50 μm radius in mammalian brain^35,54,73^. The FBR is speculated to contribute to this discrepancy^54^, as fewer viable neurons are observed surrounding electrodes after chronic implantation^74^. Particularly salient is a 60% percent reduction within 50 μm^24^. Other possible factors are that neurons remain non-active during recording sessions^54,73^ or spike-sorting methods are currently insufficient^54,70^. Presently, this discrepancy remains unresolved.

Since carbon fiber electrodes maintained a nearly natural distribution of the nearest six neuron positions, which were within 50 μm from the recording sites, we were well positioned to further consider this discrepancy by modeling electrophysiology. Also, since carbon fiber electrodes yield higher SNRs than silicon electrodes^61^ and higher SNRs are expected to yield more spikes^70^ with higher accuracy^56^, we anticipated spike sorting more clusters. Therefore, we sorted electrophysiology from an exemplary recording session towards the end of the implantation period (day 84 of 92) that yielded large spike clusters, where the mean peak-peak amplitude of the largest cluster recorded on channels that yielded clusters was 354.8±237.2 μV_pk-pk_ (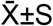, N=12 channels) and the single largest cluster had a mean amplitude of 998.9 μV_pk-pk_ (Figure 5A). Furthermore, 12 of 16 channels yielded spike clusters with mean amplitude >100 μV_pk-pk_, signifying a high chronic signal yield consistent with our previous work^62^. However, our spike sorting yielded a median of 2 and maximum of 3 spike clusters per electrode, which is considerably lower than the 18.3±4.9 neurons observed within 50 μm (N=16 fibers) even after inducing a lower FBR with high signal yield.

**Figure 5.**
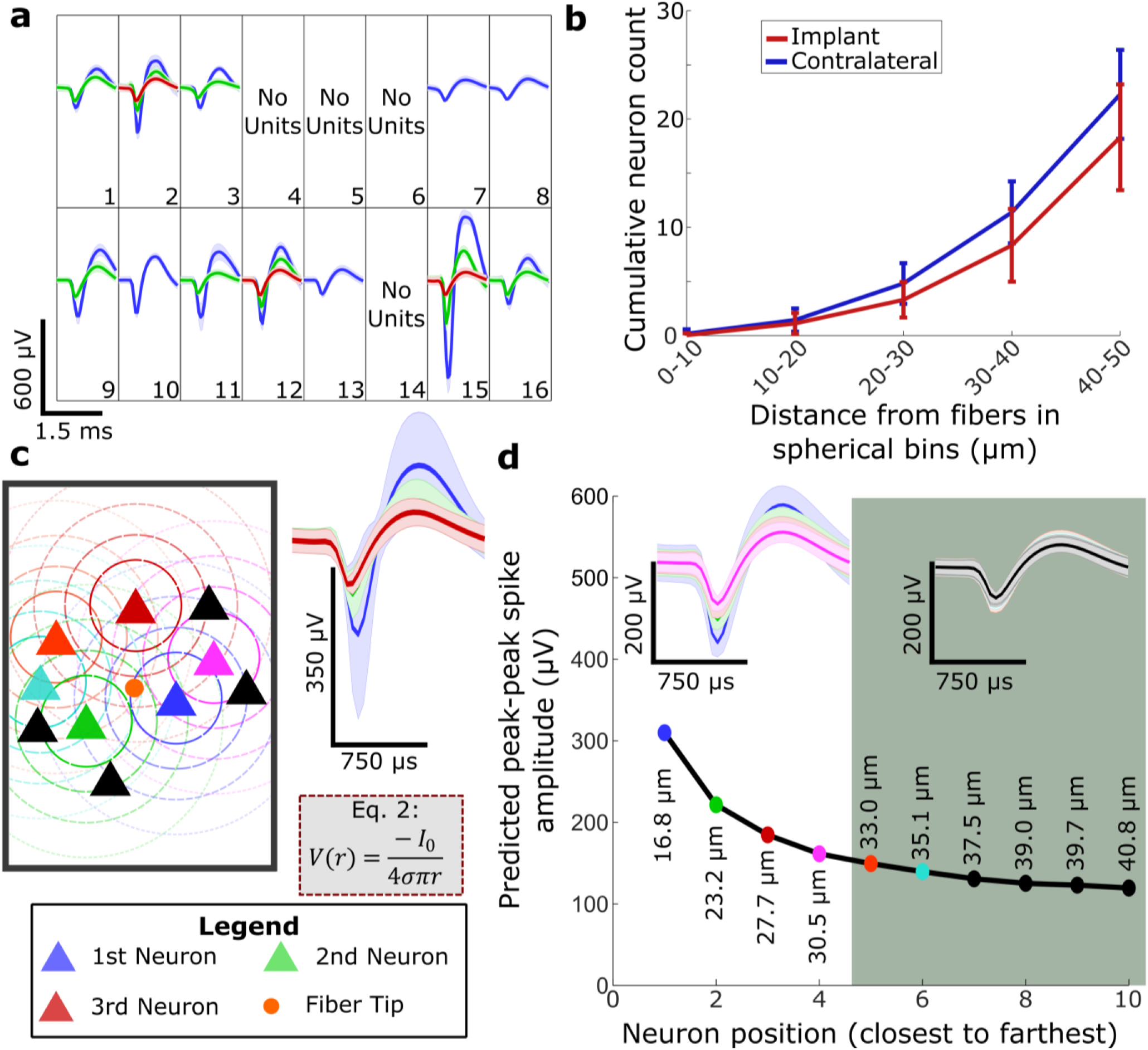
Modeling suggests neuron distribution contributes to the number of sorted spike clusters. (A) Spike sorted electrophysiology from a recording session that yielded exemplary units. Recorded on day 84 of implantation for rat #2. Solid lines show mean cluster waveform and shading shows cluster standard deviation. (B) Cumulative neuron count at each 10 μm spherical bin from implanted carbon fiber electrode tips (red). Cumulative neuron counts at each bin surrounding simulated fibers in contralateral cortex shown in blue. Error bars show standard deviation. (C) Pictorial representation of the simple point source model described by Equation 2. The carbon fiber electrode tip (orange dot) and neurons (triangles) are treated as point sources. The nearest three neurons (blue, green, and red triangles, in order) produce the three largest units (right), which were determined from all spikes attributed to the largest three units from the spike panel in (A). Neuron positions in this graphic are approximately proportional to relative distances measured in histology. Equation 2 is reproduced below the spikes. (D) Predicted mean peak-peak amplitude for the closest ten neuron positions relative to implanted carbon fibers as observed from histology, where the color of each dot corresponds to the neuron of the same position in (C). Neuron positions are in order from closest to farthest. Mean spike waveforms matching mean peak-peak amplitude are displayed above the curve and match each corresponding point by color. Waveforms are divided into two zones, the “easy sorting zone” (left, white background), where neighboring neuron spike amplitudes differ by > 15 μVpk-pk, and the “hard sorting zone” (right, grey background) where neighboring neuron spike amplitudes differ by < 15 μVpk-pk. See also Figure S5 for all waveforms grouped into one plot. Shading shows standard deviation.

We hypothesized that neuron distribution itself may contribute to the low number of sorted clusters. As expected^34^ from the geometry of concentric spheres with increasing radius, the number of neurons relative to the electrode grew rapidly with increasing spherical radius. On average, we found fewer than one neuron (0.0±0.2) within 10 μm, three neurons (3.3±1.6) within 30 μm, and fifteen neurons (15.3±4.1) 30-50 μm away from fiber tips (Figure 5B). Given the inverse relationship between neuron distance and the recorded spike amplitude^35,54,69^, the large neuron count at further distances would likely generate similar spiking amplitudes that would be difficult to sort^54^.

We used the simplest point source model where neurons are treated as points^68,75^ that have the same spiking output^46^. Figure 5C illustrates this model with the relative distances of the closest ten neurons based on our measurements (Table 2). We fit the model for *I_0_* using spikes sorted into the largest three units’ mean spike amplitudes (N=12 channels) (Figure 5C) and the nearest three neurons’ mean positions (Table 2), where conductance, *σ*, was 0.27 S/m^76^. This fit yielded *I_0_* = 16.4 nA (Figure S3A), which is similar to the 10 nA determined from fitting 60 μV to 50 μm, as recommended by Pedreira *et al*.^54^ from combined intracellular and extracellular recordings^34,35^. To verify this parameter, we fit the model in two other ways by matching each sorted cluster with a prospective neuron positioned around the same electrode (e.g., the second largest cluster associated with the second closest neuron) (Figure S3B). Fitting for both *I_0_* and *σ* yielded similar values of 14.2 nA and 0.23 S/m, respectively, while fitting for just *I_0_* yielded *I_0_* = 15.6 nA. That all three modeling approaches produced similar results (Figure S3) that were consistent with literature suggests this model can reasonably predict spike amplitudes with neuron distance as input.

Using their average relative positions to implanted fibers in the model, we plot the average spike amplitudes of the nearest ten neurons to determine spike cluster discriminability (Figure 5D). As expected from the neuron distribution we observed, predicted amplitudes quickly approached an asymptote as neuron position increased. Since baseline noise may contribute to spike detection and cluster differentiability^51,54^, we hypothesized that noise may obfuscate these small differences in amplitudes from neighboring neurons. Our previous work measured baseline noise of 10 and 15 μV_rms_ in Michigan-style silicon probes and carbon fiber electrodes, respectively^61^. Therefore, we compared the average difference in spiking amplitude between consecutive neurons to these noise levels to estimate neuron differentiability (Figure S4). This difference remained smaller than 15 μV_p-p_ for comparisons of neuron positions beyond the third and fourth neuron and smaller than 10 μV_p-p_ for comparisons beyond the fourth and fifth neuron. Therefore, the fourth closest neuron (30.5±4.5 μm) is situated along the boundary at which spike clusters become indistinguishable, capturing activity from multiple neurons, and is consistent with the 1-4 sortable clusters typically observed in literature^72^. To further illustrate, we grouped simulated spikes according to differentiability, where the four largest clusters are more easily separable than the fifth through the tenth (Figure 5D) and plotted all ten waveforms together (Figure S5). When separated, the four largest units could easily merge into two or three units with variation in position and consequently amplitude. This is plausible given that within the four closest neurons observed for 11/28 fibers, at least two neighboring neuron positions were within 1 μm apart. Similarly, when plotted together, the clusters begin to merge starting with the fourth largest cluster.

## Discussion

Here, we localized the positions of carbon fiber electrode tips that had been chronically implanted in motor cortex with subcellular-scale confidence for the first time. Volumetric confocal imaging of the implant site enabled this localization and an examination of the surrounding neurons and FBR with more 3D details and precision^20,21,23,39,59–62^. In particular, we focused on the nearest 50 μm regarded as the single-unit recording zone^34,35^, in which we observed a neuron distribution similar to that of healthy cortex. In contrast, multi-shank silicon arrays such as the UEA can remodel the nearby neuron distribution by inducing large tissue voids^20,21^ and widespread necrosis^77^. A more direct comparison can be made with Michigan-style electrodes, which can induce a 60% neuron loss within the 50 μm recording radius^24^. Given that we observed an 18% neuron loss on average within this radius of carbon fibers, a conservative estimate suggests that carbon fibers retain at least nine more neurons per penetrating electrode on average without accounting for the shanks’ larger sizes. When scaling up to the 100+ channels used in BMI devices, several hundreds of neurons could be preserved by carbon fibers instead of silicon shanks. Previously, we have also shown that our device has much higher functional probe yield and records more units per probe with better recording SNR^20,61,62^. Therefore, our work provides strong evidence that subcellular-scale electrodes such as carbon fibers retain recordable neurons to a considerably greater degree.

At the same time, our detailed examination revealed components of the FBR that could inform future designs. This included neuron soma stretching around carbon fibers, similar to that was observed surrounding silicon electrodes, which was attributed to chronic micromotion relative to surrounding tissue^28^ or to electrode insertion during surgery^29^. Also, the persistence of astrocyte enrichment along electrode tracts to depths beyond the silicon support tines suggests that permanent silicon shuttles, even with small feature size (15.5-40.5 μm), reduce biocompatibility. As sharpening the tips of electrodes with microwire-like geometry enables facile and unsupported insertion to deep cortical depths (1.2 mm^78^-1.5 mm^79^), future iterations could use fire-sharpening^78,80^ or electro-sharpening^79,81^ to obviate the need for shuttles and to reduce insertion force^79^. Furthermore, skull-fixing the electrodes induces an increased FBR compared to floating electrodes^82^. Given that the neuron distortion observed here likely accompanied implant micromotion^28^, subcellular-scale electrode designs should adopt a floating architecture similar to that of the UEA, as flexible electrodes such as syringe-injectable^39,83,84^ and NET^40^ probes already have, even with the added benefits of their small size.

Having visualized an approximately naturalistic neuron distribution around carbon fibers and recorded electrophysiology from them, we sought to characterize the relationship between the surrounding neuron population and spikes recorded through modeling. We considered predicting individual neuron positions, but tip localization would need to be more precise, as our estimated localization error (9.6 μm, Table 1) was higher than the average difference in position between the first and second closest neurons (6.4±5.4 μm) and subsequent positions (Figure S6). Regardless, modeling the ten closest neurons’ spike amplitudes suggests that the fourth closest may be situated on the boundary at which clusters become indistinguishable and may explain the low number of clusters (1-4)^72^ attributed to individual neurons. That this single unit boundary is 30.5±4.5 μm away and dependent on neuron distribution disagrees with the notion that this boundary is 50 μm^34,35^. That said, spikes from neurons up to 140 μm away have been reported as distinguishable from background noise in the hippocampus, thereby contributing to multi-units beyond the above-mentioned single unit boundary^34,35^. However, this distance may also be brain region dependent, because recording in regions with varied neuron densities or cell type distributions^85,86^ likely produce distinct background noise levels^46^ that heavily influence spike detectability. Therefore, the neuron density of target regions must be considered when interpreting intracortical electrophysiology and should influence electrode design parameters such as recording site pitch^87^. Additionally, that our exemplary recording session yielded a median of two clusters suggests that other previously identified factors, such as silent neurons^54,73^ (although disputed^50^), limitations in spike sorting algorithms^54^, and baseline noise^51^ may have contributed to the low cluster count. Furthermore, recent work suggests that favorable histological outcomes may not correlate with high recording yield^88^. This may be explained by neuronal hypoexcitability following electrode implantation^29^ or an observed shift toward a higher proportion of activity from inhibitory neurons surrounding chronic implants^26,88^. Therefore, our results contribute to an increasing list of factors that comprise the mismatch between cluster sortability and histology. Combined with recent work demonstrating that the downstream biological effects of electrode implantation are more complex than traditionally thought, our modeling and assessments of the FBR corroborate the need for further investigation into the interactions between electrodes and surrounding tissue, even for designs more biocompatible than traditional silicon electrodes^30,36,38,88^.

## Acknowledgements

We would like to thank Paul Jenkins who provided antibodies for immunohistology and Khalil Najafi for allowing use of his equipment for fabrication. We thank Chris Andrews’ help with statistics. We thank Rachel O’Farrell, Elissa Welle, Eric Kennedy, and Julianna Richie’s help with recording electrophysiology. We also thank Julianna Richie for assistance with array fabrication and with references. We also thank the staff of the Lurie Nanofabrication Facility for their expertise. This work was financially supported by the National Institute of Neurological Disorders and Stroke (UF1NS107659) and the National Science Foundation (1707316).

## Author Contributions

Conceptualization: JGL, PRP, JDW, CAC, and DC. Methodology: JGL, PRP, LAW, CAC, and DC. Software: JGL, PRP and LAW. Formal Analysis: JGL, PRP, J-CH and IMSF. Investigation: JGL, PRP & EdV. Resources: EdV, JDW, CAC, and DC. Data Curation: JGL, PRP, J-CH, and IMSF. Writing – Original Draft: JGL, CAC, and DC. Writing – Review & Editing: PRP, LAW, and JDW. Visualization: JGL, PRP, CAC, and DC. Supervision: CAC and DC. Project Administration: CAC and DC. Funding Acquisition: CAC and DC.

## Methods

### Carbon fiber electrode array fabrication

High Density Carbon Fiber (HDCF) electrode arrays with 16 channels were fabricated using methods reported elsewhere^63^. Briefly, silicon support tines were fabricated from 4” silicon wafers using silicon micromachining processes. The support tines had trenches etched into them via deep reactive ion etching to hold the fibers for facile insertion into the brain, tapered to a width of 15.5 μm, had a pitch of 80 μm, and had a length of 3 mm for targeting cortex. The support tines had gold pads on them to interface with the carbon fibers. These gold pads then led to a separate set of gold bond pads to interface with a printed circuit board (PCB). Once tines were fabricated, they were bonded to a custom PCB with Epo-Tek 301 epoxy (Epoxy Technology, Billerica, MA). 2-Part epoxy (1FBG8, Grainger, Lake Forest, IL) was applied to the underside of the silicon portion to provide buttress support. The gold bond pads were then wire-bonded to pads on the PCB and the wire bonds coated in Epo-Tek 353ND-T epoxy. Carbon fibers were then laid into the support tines by exploiting capillary action from a combination of deionized water (electrical pad end) and Norland Optical Adhesive 61 (NOA 61) (Norland Products, Inc., Cranbury, NJ) (distal end). A NLP 2000 (Advanced Creative Solutions, Carlsbad, CA) was used to apply Epo-Tek H20E silver epoxy to the gold pads and the carbon fibers to establish an electrical connection. NOA 61 was applied to the gold pads and the carbon fibers to further secure them. Fibers were then cut to a length of 1000 μm and coated with ~800 nm of Parylene C (PDS2035CR, Specialty Coatings Systems, Indianapolis, IN). Fibers were then laser cut to a final length of 300 μm beyond the silicon support tine ends with a 532 nm Nd:YAG pulsed laser (LCS-1, New Wave Research, Fremont, CA) as described previously^62^. Carbon fibers were then plasma ashed in a Glen 1000P Plasma Cleaner (Glen Technologies Inc., Fremont, CA). Fiber tips were functionalized in one of two ways: 1) electrodeposition by dipping carbon fibers in a solution of 0.01 M 3,4-ethylenedioxythiophene (483028, MilliporeSigma, Darmstadt, Germany) and 0.1 M sodium p-toluenesulfonate (152536, MilliporeSigma) and applying 600 pA/channel using a PGSTAT12 potentiostat (EcoChemie, Utrecht, Netherlands) to coat the tips with PEDOT:pTS^25,61^ (N=3 arrays) or 2) electrodeposition of Platinum Iridium (PtIr) with a Gamry 600+ potentiostat (Gamry Instruments, Warminster, PA) (N=2 arrays) using previously published methods^89^.Silver ground and reference wires (AGT05100, World Precision Instruments, Sarasota, FL) were soldered to the PCB, completing assembly. For one electrode, a support tine was broken off prior to implant as a sham channel for a separate study (for rat #1).

### Electrode implantation

All animal procedures were approved by the University of Michigan Institutional Animal Care and Use Committee

Adult male Long-Evans rats (N=5) weighing 393-630 g were implanted with one HDCF electrode array each. Surgical implantation closely followed procedures performed in our previous work^62,90^. Throughout the surgeries, temperature was monitored with a rectal thermometer and breath rate was monitored with a pulse oximeter. Isoflurane (5% (v/v) induction, 1-3 % maintenance) was used as a general anesthetic and carprofen (5mg/kg) as a general analgesic. After opening the scalp, 7 bone screws (19010-00, Fine Science Tools, Foster City, CA) were screwed into the skull. One screw at the posterior end of the skull was used for referencing. A 2 × 2 mm craniotomy was drilled in the right hemisphere, where the bottom left corner of the craniotomy was 1 mm lateral and 1 mm anterior to bregma. The probe was then lowered to the dura mater to zero its dorsal/ventral position. After durotomy with a 23G needle, the probe was immediately inserted to a depth of 1.2-1.3 mm to reach layer V of motor cortex. The craniotomy was then filled with DOWSIL silicone gel (DOWSIL 3-4680, Dow Silicones Corporation, Midland, MI). Ground and reference wires were wrapped around the most posterior bone screw for referencing. A headcap was formed by applying methyl methacrylate (Teets denture material, 525000 & 52600, Co-oral-ite Dental MFG. Co., Diamond Springs, CA) onto the skull until the probe’s electrical connector was firmly in place and bone screws were covered. The scalp was sutured around the connector and surgery was complete.

### Electrophysiological recording and spike sorting

Electrophysiological recordings were collected in chronically implanted rats while awake and freely moving in a Faraday cage as we have reported previously^62^. Signals were recorded using ZC16 and ZC32 headstages, RA16PA pre-amplifiers, and a RX7 Pentusa base station (Tucker-Davis Technologies, Alachua, FL) at 24414.1 Hz. Recordings were collected at least weekly for 10 minute sessions. Spike sorting was semi-automated and based upon procedures we have described previously^61,62^. Sessions collected towards the end of the implantation period were chosen to increase coherence with histological outcomes^88^. Channels were excluded from a session if the impedance at 10 Hz was abnormally high compared to other channels and previous sessions (~1-2 weeks), which was similar to our exclusion criteria in previous work^61^. For the exclusion determination, impedance was measured using electrochemical impedance spectroscopy as we have described previously^61^. Common average referencing was performed using the remaining channels to reduce noise^91^. The following steps were performed in Plexon Offline Sorter (version 3.3.5) (Plexon Inc., Dallas, TX) by a trained operator. Signals were high-pass filtered using a 250 Hz 4-pole Butterworth filter. Five 100 ms snippets of signal with low neural activity and artifact noise were manually selected from each channel and used to measure V_RMS_ noise for each channel^61^. The threshold for each channel was set at −3.5*V_RMS_. Cross channel artifacts were then invalidated. Putative clusters centers were manually designated and waveforms assigned using K-Means clustering. Obvious noise waveform clusters were removed. Automated clustering was performed using the Standard Expectation-Maximization Scan function in Plexon Offline Sorter^62^. Persisting noise waveforms were removed, and obvious oversorting and undersorting errors were manually corrected. Clusters were also cleaned manually. Resultant waveforms were imported into and analyzed in MATLAB (version R202b) (MathWorks, Natick, MA) using custom scripts.

### Tissue preparation, immunohistochemistry, and imaging

At the end of the implantation period, rat brains were prepared for immunohistochemistry and histological imaging. Rats were transcardially perfused on day 88-92 (N=3) or day 42 (N=2) as described previously^61,62^. If the perfusion fixation was successful, brains were extracted and soaked in 4% paraformaldehyde (PFA) (19210, Electron Microscope Sciences, Hatfield, PA) in 1x phosphate buffered saline (PBS) for 1-3 days. If more fixation was required, brains remained in the skull while soaking in PFA solution for two days before extraction, followed by an additional 24-hour incubation in PFA solution. In all cases, the electrode array, headcap, and skull-mounted bone screws were removed from the brain. Brains were then incubated in 30% sucrose (S0389, MilliporeSigma) in 1x PBS with 0.02% sodium azide (S2002, MilliporeSigma) for at least 72 hours until cryoprotected. Brains were then sliced to a thickness of 300 μm^59^ with a cryostat. Slices were selected for staining based upon estimated depth and/or from the observation of holes in a brightfield microscope.

Immunohistochemistry closely followed staining techniques we have used previously^62^, but modified to accommodate 300 μm brain slices^59^. All incubation periods and washes were performed with brain slices in well plates on nutators. Chosen slices were first incubated in 4% PFA (sc-281692, Santa Cruz Biotechnology, Dallas TX) for 1 day at 4°C. Slices were washed for 1 hour in 1x PBS twice at room temperature and then incubated in a solution containing 1% Triton X-100 (93443, MilliporeSigma) in StartingBlock (PBS) Blocking Buffer (37538, ThermoFisher Scientific, Waltham, MA) overnight at room temperature to permeabilize and block the tissue, respectively. Slices were then washed in 0.1-0.5% Triton X-100 in 1x PBS (PBST) solution for one hour at room temperature three times.

Slices were incubated in primary antibodies for 7 days at 4°C, where antibodies were added to a solution containing 1% Triton X-100, 0.02% sodium azide in 1x PBS, and StartingBlock. The primary antibody cocktail differed between rats implanted for 6 weeks (N=2) and 12+ weeks (N=3). In all rats, antibodies staining for neurons (Mouse anti-NeuN, MAB377, MiliporeSigma) and astrocytes (Rabbit anti-GFAP, Z0334, Dako/Agilent, Santa Clara, CA) were used, where both had dilution ratios of 1:250. For 6 week rats, staining for axon initial segments (AIS) with Goat anti-Ankyrin-G (1:1000 dilution ratio) was added. The ankyrin-G antibody was made and provided by the Paul Jenkins Laboratory (University of Michigan, Ann Arbor, MI) using methods published previously^92^. For 12+ week rats, staining for microglia (Guinea Pig anti-IBA1, 234004, Synaptic Systems, Göttingen, Germany) (1:250 dilution ratio) was added. Slices were washed in 0.1-0.5% PBST three times at room temperature for one hour each wash before secondary antibody incubation at 4°C for 5 days. Secondary antibodies used the same base solution as the primary cocktail. The following secondary antibodies were used: Donkey anti-Mouse Alexa Fluor 647 (715-605-150, Jackson ImmunoResearch Laboratories, Inc., West Grove, PA) for neurons, Donkey anti-Rabbit Alexa Fluor 546 (A10040, Invitrogen, Carlsbad, CA) for astrocytes in 6 week rats, Goat anti-Rabbit Alexa Fluor 532 (A11009, Invitrogen) for astrocytes in 12+ week rats, Donkey anti-Guinea Pig Alexa Fluor 488 (706-545-148, Jackson ImmunoResearch Laboratories, Inc.) for microglia, Donkey anti-Goat-Alexa Fluor 488 (705-545-003, Jackson ImmunoResearch Laboratories, Inc.) for AISs, which all had dilution ratios of 1:250, and 4’,6-Diamidino-2-Phenylindole, Dihydrochloride (DAPI) (D1306, Invitrogen) for cellular nuclei (1:250-1:500 dilution ratio). Slices were washed in 0.1-0.5% PBST twice at room temperature for two hours each wash and were washed in 1x PBS with 0.02% azide for at least overnight before imaging could commence. Stained slices were stored in 1x PBS with 0.02% azide at 4°C outside of imaging sessions after staining.

Prior to imaging, slices were rapidly cleared using an ultrafast optical clearing method (FOCM)^93^. FOCM was prepared as a solution of 30% (w/v) urea (BP169500, ThermoFisher Scientific), 20% (w/v) D-sorbitol (DSS23080-500, Dot Scientific, Burton, MI), and 5% glycerol (w/v) (BP229-1, ThermoFisher Scientific) in dimethyl sulfoxide (DMSO) (D128-500, ThermoFisher Scientific). Glycerol was added at least one day after other reagents started mixing and after urea and D-sorbitol were sufficiently dissolved in DMSO. FOCM was diluted in either MilliQ water (N=1 rat) or 1x PBS (N=4 rats) to 25%, 50% and 75% (v/v) solutions. During clearing, slices were titrated to 75% FOCM in 25% concentration increments, 5 minutes per step. 75% FOCM in water was found to expand tissue laterally (6.7%) after imaging rat #2, so other slices from other rats were cleared with FOCM in 1x PBS. This expansion was not corrected in subsequent analyses. The clearing process was repeated immediately before each imaging session. The samples remained suspended in the 75% FOCM solution during imaging. Images were collected with a Zeiss LSM 780 Confocal Microscope (Carl Zeiss AG, Oberkochen, Germany) with 10x and 20x objectives. The microscope recorded transmitted light and excitation from the following lasers: 405, 488, 543, and 633 nm. Z-stacks of regions of interested were collected along most, if not all, of the thickness of the sample. Images had XY pixel size of 0.81 μm or 0.202 μm and z-step of 3 μm or 0.5-0.6 μm, respectively. Laser power and gain were adjusted to yield the highest signal-to-noise (SNR) ratio while also minimizing photobleaching. The “Auto Z Brightness Correction” feature in ZEN Black (Carl Zeiss) was used to account for differences in staining brightness along the slice thickness. The implant site was located by overview imaging and quick scanning, where high astrocyte staining signal in dorsal regions and a straight line of holes delimited the electrode site (Figure 1B). Sites with approximately similar stereotaxic coordinates to those of the implant site but in the contralateral hemisphere of the same slice were imaged as control. After imaging, slices were titrated back to 0.02% sodium azide in 1x PBS, and stored. Clearing was repeated again when further imaging was needed, as each implant site required multiple imaging sessions.

### Putative tip localization

Putative carbon fiber electrode tips were localized manually by three reviewers experienced in histology. Tips were localized using ImageJ (Fiji distribution, multiple versions)^94^ with additional cross-referencing in Zen Black (2012) (Carl Zeiss). Fiber tips were localized by first identifying electrode tracts in more dorsal focal planes. These tracts presented as dark holes in transmitted light and fluorescent imaging channels that were typically positioned in an approximately straight line and surrounded by high GFAP intensities. Putative electrode tracts were more easily visualized by using contrast adjustment, histogram matching^66^ to account for changes in brightness throughout the brain slice, median or Gaussian filters (including 3D versions^67^) for reducing noise, maximum intensity projections, and toggling the imaging channels that were visualized simultaneously. Tips were then located by scrolling through z-stacks toward ventral focal planes. The estimated tip depth was determined by finding the z-step at which the tract rapidly began to reduce in size and become filled with surrounding background fluorescence or parenchyma when scrolling dorsal-ventrally through the image. The tip width, height, and center were determined by manually fitting an ellipse to the electrode tract at that z-step.

### Verification of tip localization

Putative tip localization was verified by measuring electrode tract pitches, tip cross-sectional area, and the agreement between three reviewers in tip localization. Electrode tract pitches were measured in ImageJ after stitching images^65^. The positions of electrode tracts were measured in the same z-step or in nearby z-steps to achieve an approximately planar arrangement of tracts. Ellipses were manually fit to tracts, and the Euclidean distance between the centers of neighboring tracts were measured in MATLAB to determine pitch. The cross-sectional area of tips was calculated as the area of an ellipse fit to fiber tips localized as described above. The agreement between three reviewers experienced in histology was determined by measuring the absolute value of the Euclidean distance between the three centers of the ellipses fit to each fiber. When determining the localization confidence, all comparisons for each group of fibers (see Table 1) were grouped together. The median and interquartile range were measured from each group of comparisons for Table 1.

Fibers tips that were not localized by all three reviewers were excluded from further analyses and from the agreement determination in Table 1. Three fibers were excluded from rat #5 because one fiber was not found by one reviewer, and two fibers were found in different images than the other reviewers, so localizations were not directly comparable. Tissue damage in rat #3 was so severe (Figure S1) that even the number of fibers in the tissue was not consistently identified by reviewers, therefore rat #3 was excluded from quantitative analyses. For cross-sectional area measurements and subsequent analyses requiring fiber localization, the positions localized by the first author were used, where exclusion from other reviewers were not considered.

### Segmenting neuron somas into 3D scaffolds

The somas of neurons surrounding electrodes were manually segmented in 3D after putative fiber localization to measure soma morphology and determine the positions of neuron centroids. Image pre-processing and segmentation was performed using ImageJ plugins. The NeuN channel was used to visualize neuron somas for segmentation. First, a z-step with a qualitatively high SNR was selected and contrast adjusted to make the image clearer. The remaining z-steps in the z-stack were histogram matched^66^ to the selected z-step to account for variations in brightness throughout the slice along its thickness. A 3D median filter^67^ was applied to reduce noise. The ImageJ plugin nTracer (primarily version 1.3.5)^95^ was used to segment each neuron by manually tracing along the cell outline in each z-step that the neuron was present (Figure 2E). Once complete, a 3D scaffold of neuron somas was produced (Figures 2F-G). Neurons that bounded voxels that were within a 100 × 100 μm (diameter X height) cylinder centered at the electrode tip locations were selected for tracing. Neurons were also traced in contralateral regions as a control. For rat #1, five points with similar coordinates to fiber tip centroids were selected and neurons traced within a cylinder (100 μm diameter, 100 μm height) (N = 5 hypothetical fibers, N=199 neurons). For rat #2, all neurons that bounded voxels in a cylindrical region (300 μm diameter, 300 μm height) were traced (N=926 neurons). Traced neurons were imported into MATLAB using a custom script.

For some fibers (rat # 1: N=8 fibers; rat #2: N=5 fibers), the electrode tip was closer to the top of the brain section than 50 μm. Instead of 50 μm, the distance between the fiber tip and the top of the brain section at that fiber’s tract was selected as half the cylinder height for tracing and used in subsequent analyses as the radius around that fiber. Additionally, neuron somas that were cut along the plane of cryosectioning were excluded if it was clear that the middle of the soma was excluded from the brain slice.

### Neuron morphometrics and densities

Neuron morphometrics and positions were extracted from the 3D scaffolds of neurons produced by tracing. To measure neuron soma volume, all voxels bounded by the scaffold were counted and the total number was multiplied by the volume of a single voxel^96^. The centroids of these sets of points were used to determine neuron positions and distances relative to the putative electrode tips. Centroid positions were also used for counting the number of neurons within 3D radii, such as 50 μm, and in determining neuron density. To determine the extent of neuron elongation around implanted fibers, the cell shape strain index (CSSI) as defined by Du *et al*. with Equation 1^28^ was measured. An ellipse was fit to the soma trace in the z-step that had the greatest cross-sectional area for that neuron (Ohad Gal. fit_ellipse [https://www.mathworks.com/matlabcentral/fileexchange/3215-fit_ellipse], MATLAB Central File Exchange). CSSI was determined using Equation 1^28^, where *a* is the minor axis and *b* is the major axis from elliptical fitting.

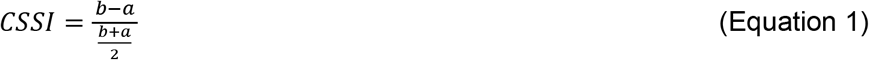

These same measurements were repeated for neurons in contralateral sites. Comparisons of neurons within a 50 μm spherical radius of localized tips (Figures 3C and 3E) included neurons where the centroid was within 50 μm. To determine the relationship between distance and CSSI or volume (Figures 3D and 3F), only neurons that were ventral to all tips analyzed in the source image were included to remove the effects of multiple nearby tips.

### Nearest Neuron Positions

Neuron positions relative to implanted fibers were determined by calculating the Euclidean distance between the center of the ellipse manually fitted to each carbon fiber electrode tip (see *Tip localization*) and centroid of all points bounded by each traced neuron. These distances were sorted from shortest to longest to determine the nearest neuron positions relative to implanted fibers. When determining the mean neuron position, if that position was not measured for one or more fibers due to a reduced tracing radius, that fiber was excluded from the mean neuron position calculation for that position (see Table 2).

Similar measurements were performed at contralateral sites as a control. In N=1 rat, neurons within 50 μm of five hypothetical fiber points with similar coordinates to localized fiber tips were traced, and the nearest neuron positions were determined in the same manner as with implanted fibers. For another rat, all neurons within a 300 × 300 μm (diameter X height) cylindrical volume were traced. Points were placed in a 3D grid with 12.5 μm spacing within this volume but were excluded if positioned within 50 μm of the volume’s border or if within the set of points bounded by a traced neuron. These points were used as hypothetical carbon fiber tip locations in unimplanted contralateral tissue. Similarly to implanted fibers, the Euclidean distance between each hypothetical point and the centroid of all points bounded by each traced neuron was measured to determine relative neuron distances. As described earlier (see *Neuron morphometrics and densities*), these neuron centroid positions were also used to determine the neuron densities and neuron counts within spherical radii from hypothetical fiber locations.

### Glial response around implanted carbon fibers

Glial responses to chronic implantation of carbon fiber electrodes were evaluated by measuring the staining intensity of GFAP and IBA1 surrounding implanted fibers, similar to measurements commonly made previously (e.g., ref^61^ and ref^97^). Z-steps that captured focal planes that included regions of missing tissue due to capturing either the top or bottom of the brain slice were excluded. Fibers were excluded if their tips were localized in excluded z-steps. Confocal images were imported into MATLAB using the Bio-Formats MATLAB Toolbox (The Open Microscopy Environment, https://www.openmicroscopy.org/bio-formats/downloads/) for analysis. The background intensity for each z-step was determined first. The Euclidean distance of each pixel in the z-step to every tip center was determined. If the z-step was dorsal to the tip, then the distance to the center of the electrode tract in that z-step was determined instead. The mean intensity of pixels that were 300-310 μm away from those tip positions was measured as the background intensity for that z-step. Next, a line (the electrode axis) was fit to the coplanar fiber tip positions. The minimum pitch of the electrode tip locations along this line was used to determine bounded lanes centered at each tip for measurement. The intensity of glial fluorescence at increasing distances from each electrode tip were measured by determining the mean intensity of pixels bounded by concentric rings that were 10 μm thick, centered at each electrode tip location, and were bounded by these lanes with equal width to prevent overlap between fibers. Glial intensity was reported as the ratio of the mean intensity for each bin to the mean intensity of the background for the z-step containing the tip. Since fiber tips were localized at a range of z-steps, the measurement was performed at the appropriate z-step for each fiber.

This process was performed for both GFAP and IBA1. As an additional control, the process was repeated using images collected of sites in the same brain sections as the implanted tips with similar stereotaxic coordinates in the contralateral hemisphere.

### Modeling of Electrophysiology

A simple point source model^68^, similar to what we have used previously^75^, was used to predict the extracellular spike waveforms recorded by carbon fiber electrodes that originated from neurons surrounding implanted carbon fibers. This model is defined by Equation 2:

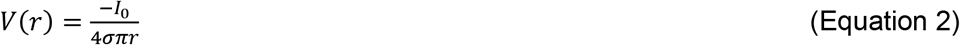

where *V* is the extracellular potential recorded by an electrode, *I_0_* is the extracellular current generated by a neuron firing an action potential, *σ* is the conductivity of the brain tissue between the electrode and the neuron, and *r* is the relative distance between the neuron and the electrode. The model operates on the following assumptions: 1) that the brain is an isotropic medium with regard to frequency^98^, 2) neurons are treated as point sources^68,75,99^ 3) cell spiking output over time is constant^69^ and across neurons^46,75^. Using values for *I_0_* and *σ* that were empirically fit or identified in literature renders the equation a single-parameter model, where the relative distance of the source neuron to the electrode, *r*, is the required input.

The model was fit empirically using three methods. In the first, *σ* = 0.27 S/m was selected from literature^76^ and *I_0_* was determined from fitting. Spike clusters that were recorded 84 days post implant in rat #2 were sorted (Figure 5A) and ranked by mean amplitude in descending order. Spikes associated with the largest cluster recorded on each channel that recorded sortable units (N=12 channels) were grouped, and the mean peak-peak amplitude was plotted against the mean position of the closest neuron (N=2 rats). This was repeated for the second and third largest clusters and matched to the second and third closest neuron positions from Table 2, respectively. These waveforms are shown in Figure 5C (right). *I_0_* was then fit to these three value pairs using the MATLAB function *lsqcurvefit* (Figure S3A). In the second and third methods, the fit was performed using individual spike cluster and neuron distance pairs (Figure S3B). For each channel, spike clusters were sorted in descending order by mean peak-peak waveform amplitude. These cluster amplitudes were plotted against the positions of neurons surrounding those channels with the same rank in position (e.g. the amplitude of the second largest cluster plotted against the position of the second closest neuron). One cluster was excluded because the position of the third closest neuron was not measured due to the electrode tip’s proximity to the top of the brain slice. In the second method, both *σ* and *I_0_* were fit using *lsqcurvefit*. In the third method, *σ* = 0.27 S/m and *I_0_* was fit.

To plot predicted waveforms, all spikes sorted across all clusters on day 84 for rat #2 (Figure 5A) were collected and normalized to have peak-peak amplitudes of 1. The mean positions of the nearest ten neurons were used as input into Equation 2 using the first empirical fit, which yielded the scaling factor for the normalized spikes. The mean and standard deviation of the scaled spikes was evaluated for each position and plotted (Figures 5D and S5).

### Statistical Analyses

Comparisons of neuron soma volumes and CSSIs between the neurons surrounding implanted fibers and control regions were performed using two-sided Kolmogorov-Smirnov tests^28^ with an alpha of 0.05. Neuron densities, neuron positions, and glial intensities in radial bins surrounding the implanted fibers compared to control were performed using two-sided two sample t-tests with an alpha of 0.05. Linear regressions were determined using the *fitlm* function in MATLAB. All statistical tests were performed in MATLAB.

### Figure and graphics

All figures were generated using a combination of ImageJ, MATLAB, Inkscape (version 1.1.2), and Adobe Illustrator 2022 (version 26.1). ImageJ was used for figures showing histology. MATLAB produced numerical plots. Inkscape was used to compile and complete these figures, with some help from Adobe Illustrator.

### MATLAB code add-ons

In addition to previously stated code add-ons for MATLAB, we used the following: shadederrorbar (Rob Campbell. raacampbell/shadedErrorBar (https://github.com/raacampbell/shadedErrorBar), GitHub) (multiple versions) and subtightplot (Felipe G. Nievinski. subtightplot (https://www.mathworks.com/matlabcentral/fileexchange/39664-subtightplot), MATLAB Central File Exchange.). Both were used for figure generation.

## Video Legends

**Video 1. Volumetric imaging of implant sites captures the electrode tracts in 3D.** Volumetric imaging of the implant site along the array in rat #1 shows how the fiber tracts disappear when scrolling through the 3D image ventrally along the dorsoventral axis. The snapshots shown in Figure 1C, are taken from the same image stack. Videos start at the dorsal side of the brain slice and end approximately 30 μm ventral to the most ventral fiber tip position. The z-step resolution is 3 μm. The number in the bottom left is the distance from the first z-step, which is approximately at the top of the brain slice. Since the framerate is 2 frames (z-steps) per second, the video speed is 6 μm/s. Each segment shows a different stain in the following order: NeuN, GFAP, IBA1, DAPI, transmitted light, composite of NeuN, GFAP, IBA1 & DAPI. The images were stitched^65^, one z-step contrast adjusted, and histogram matched^66^ to that z-step in ImageJ.

**Video 2. Volumetric imaging of a single fiber tract enables tip localization in 3D.** Volumetric imaging of fiber tracts in high resolution enables localization of putative tips in 3D. This video shows the fiber tract of fiber #9 in rat #2, which is shown in snapshots in Figure 2B. The video scrolls from the dorsal side of the brain slice until the z-step 12 μm ventral to the putative tip. The number in the top left is the distance from the first z-step, which is approximately at the top of the brain slice. The tip is at the 83.4 μm mark. Top row: NeuN, GFAP and IBA1 (left to right). Bottom row: DAPI, transmitted light, and a composite of NeuN, GFAP, IBA1 and DAPI (left to right) The video has a frame rate of 5 0.6 μm thick z-steps per second, so 3 μm/s. The images were contrast adjusted in one z-step and histogram matched^66^ to that z-step in ImageJ.

