## Supplementary Material for "Profiling neurons surrounding subcellular-scale carbon fiber electrode tracts enables modeling recorded signals"

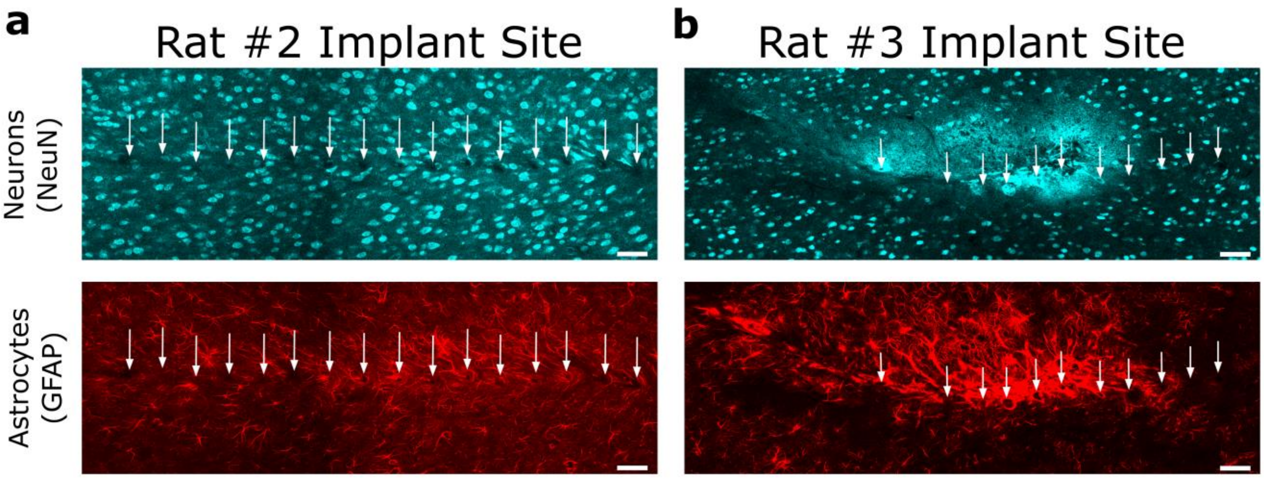


**Figure S1. Foreign body responses induced by entire carbon fiber electrode arrays chronically implanted in motor cortex were variable**. (A) Confocal images showing histology of the implant site for rat #2 showing a minimal foreign body response (FBR). The imaging plane was estimated to be up to 42 µm ventral of some tips, and up to 27 µm dorsal to others. (B) Histology of the implant site for rat #3, with a markedly greater FBR. (A and B) Scale bar, 80 µm. Cyan: NeuN; red: GFAP. White arrows point to fiber tracts identified in the tissue. Images were stitched[1] and contrast adjusted using ImageJ.

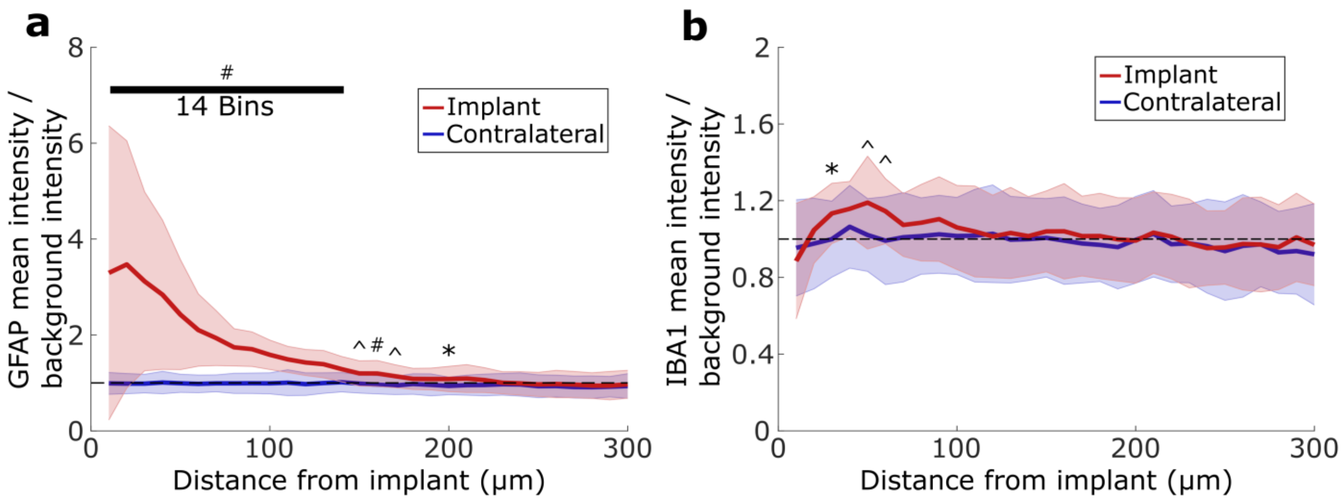


**Figure S2. Measurement of glial responses surrounding carbon fiber electrode tips.** (A) Ratio of mean GFAP intensity to background intensity over distance in 10 µm radial bins from fiber tips (N=2 rats, 26 fibers) vs. hypothetical tips (N=2 rats, N=26 fibers) in contralateral sites. Background was the mean intensity of pixels 300-310 µm from fiber locations. (B) Same as (A), but with IBA1. (A and B) implant site (red) and contralateral site (blue). * $p<0.05$. ^ $p<0.01$. # $p<0.001$. Error shading shows standard deviation.

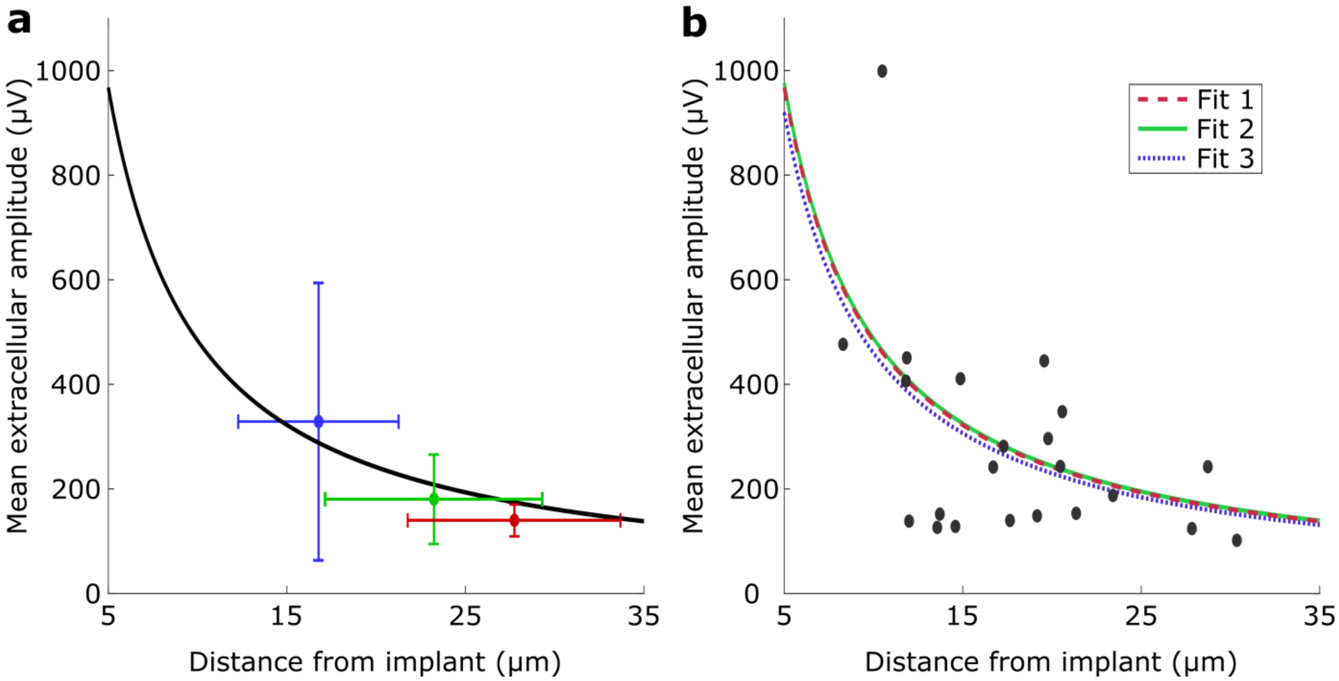


**Figure S3. Empirical fitting of the point source model.** (A) The point source model defined by Equation 2 was fit using the mean positions of the nearest three neurons and mean amplitudes of the spikes sorted with the largest three spike clusters spike sorted across all fibers in Figure 5A. The colors of the points match their respective positions in Figure 5C (Blue: first closest neuron and largest spike cluster. Green: second closest neuron and second largest cluster. Red: third closest neuron and third largest cluster). Error bars show the standard deviation of the neuron position (left to right) and standard deviation of the spike amplitudes grouped into each cluster (up and down). The black line shows the fitted model. The value for $I_0$ was fit using MATLAB's *lsqcurvefit* function. (B) The model was verified by repeating the fitting of the model using the mean amplitudes of the clusters sorted in Figure 5A and the distance observed surrounding the same channels in histology (black dots). Fit 1 is the same fit shown in (A) In Fit 2, both $\sigma$ and $I_0$ were fit. In Fit 3, only $I_0$ was fit.

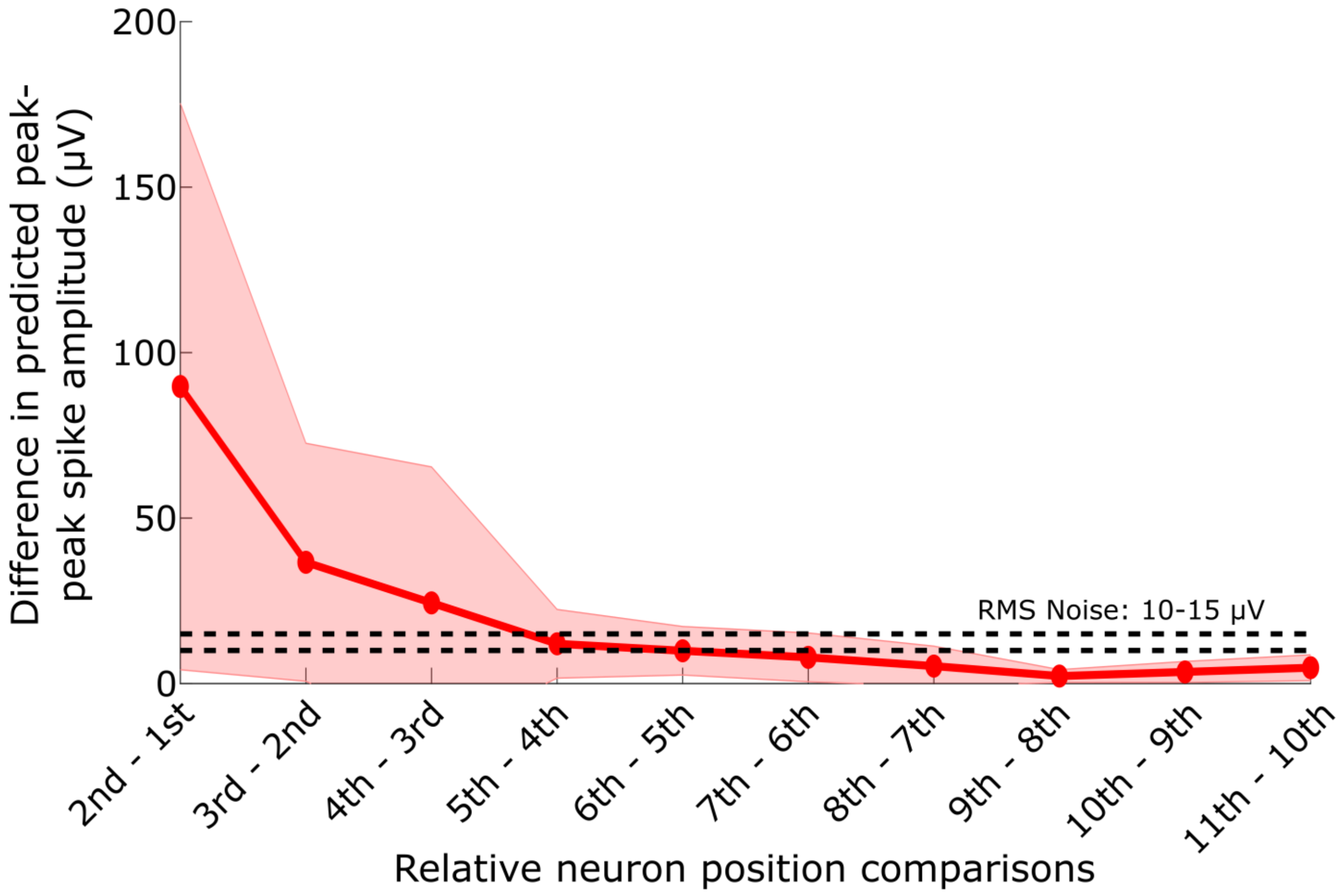


**Figure S4. Differences in predicted peak-peak spike amplitude become smaller than baseline noise.** Each point represents the average difference in average spike amplitudes of neurons of neighboring positions relative to implanted carbon fiber electrodes. The x-axis lists the neuron positions associated with each comparison, while the y-axis is the mean differential peak-peak spike amplitude resulting from that comparison. Dashed lines show baseline ($V_{rms}$) noise measured in carbon fiber electrodes (15 μV, top) and silicon shank electrodes (10 μV, bottom) reported previously[61]. Shaded error bars show standard deviation.

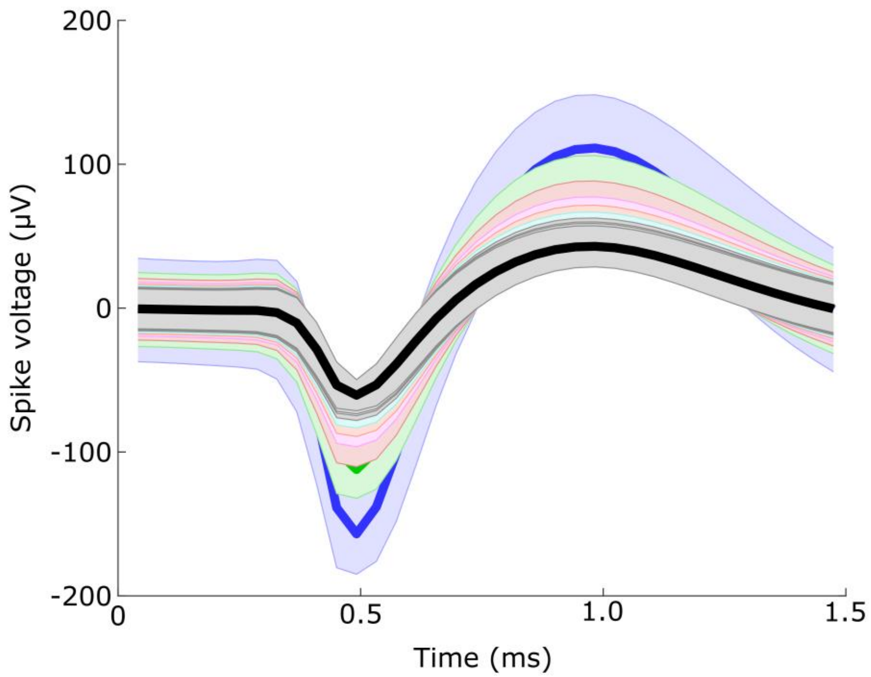


**Figure S5. Predicted mean spike waveforms for the nearest ten neurons.** The waveforms shown here are the predicted waveforms for the nearest ten neurons. These are the same waveforms shown in Figure 5D, but larger and shown in one plot to show the reduction in spike sortability starting with the fourth closest neuron in a different viewpoint.

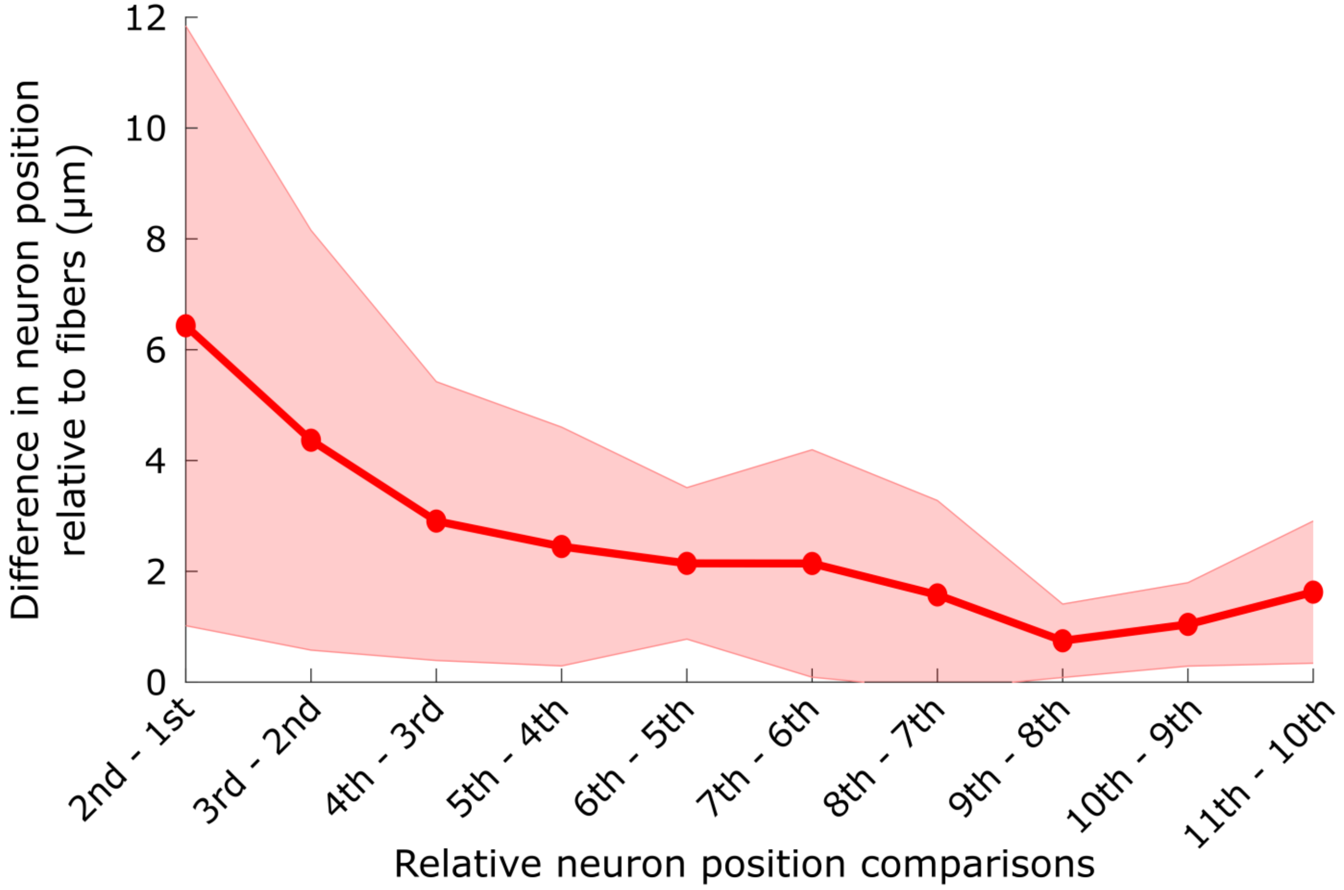


**Figure S6. Differences in relative neuron position to fibers is small.** Each point represents the average difference in average neighboring positions of neurons relative to implanted carbon fiber electrodes. The x-axis lists the neuron positions associated with each comparison, while the y-axis is the mean difference in relative position resulting from that comparison. Shaded error bars show standard deviation.

**Video 1. Volumetric imaging of implant sites captures the electrode tracts in 3D.** Volumetric imaging of the implant site along the array in rat #1 shows how the fiber tracts disappear when scrolling through the 3D image ventrally along the dorsoventral axis. The snapshots shown in Figure 1C, are taken from the same image stack. Videos start at the dorsal side of the brain slice and end approximately 30 µm ventral to the most ventral fiber tip position. The z-step resolution is 3 µm. The number in the bottom left is the distance from the first z-step, which is approximately at the top of the brain slice. Since the framerate is 2 frames (z-steps) per second, the video speed is 6 µm/s. Each segment shows a different stain in the following order: NeuN, GFAP, IBA1, DAPI, transmitted light, composite of NeuN, GFAP, IBA1 & DAPI. The images were stitched[86], one z-step contrast adjusted, and histogram matched[87] to that z-step in ImageJ.

**Video 2. Volumetric imaging of a single fiber tract enables tip localization in 3D.** Volumetric imaging of fiber tracts in high resolution enables localization of putative tips in 3D. This video shows the fiber tract of fiber #9 in rat #2, which is shown in snapshots in Figure 2B. The video scrolls from the dorsal side of the brain slice until the z-step 12 µm ventral to the putative tip. The number in the top left is the distance from the first z-step, which is approximately at the top of the brain slice. The tip is at the 83.4 µm mark. Top row: NeuN, GFAP and IBA1 (left to right). Bottom row: DAPI, transmitted light, and a composite of NeuN, GFAP, IBA1 and DAPI (left to right) The video has a frame rate of 5 0.6 µm thick z-steps per second, so 3 µm/s. The images were contrast adjusted in one z-step and histogram matched[87] to that z-step in ImageJ.
